# Analysis of hormonal regulation of promoter activities of *Cannabis sativa* prenyltransferase 1 and 4 and salicylic acid mediated regulation of cannabinoid biosynthesis

**DOI:** 10.1101/2022.11.25.517997

**Authors:** Lauren B. Sands, Samuel R. Haiden, Yi Ma, Gerald A. Berkowitz

**Author notes:** Corresponding authors: Yi Ma,; Gerald A. Berkowitz,; Agricultural Biotechnology Laboratory, 1390 Storrs Rd. Unit 4163, University of Connecticut, Storrs, CT, 06269-4163.

## Abstract

*Cannabis sativa* prenyltransferase 4 (CsPT4) and prenyltransferase 1 (CsPT1) have been shown to catalyze the step in the cannabinoid biosynthetic pathway that generates cannabigerolic acid (CBGA), the substrate for the end-point enzymes that generate cannabidiolic acid (CBDA) and tetrahydrocannabinolic acid (THCA). Prior studies from our lab suggest that CBGA production rate-limits the pathway. There is a lack of understanding concerning how important cannabinoid biosynthetic genes are regulated as cannabinoid synthesis increases during female flower development. Both *CsPT* genes were shown to be highly expressed in flowers. The genes were also found to be present in leaves and roots. GUS staining also detected the promoter activities in leaves of seedlings, and the promoter activities were drastically stronger in the section of the sugar leaves where glandular trichomes are formed. *In silico* analysis of the two *CsPT* genes revealed several hormone and transcription factor responsive elements. Dual luciferase assays were conducted to determine whether a hormone could alter the promoter activities of *CsPT1* and *CsPT4*. The results showed that *CsPT4* pro was activated following treatment from salicylic acid (SA), gibberellic acid (GA), ethylene, ABA, and cytokinin, while the *CsPT1* promoter was activated following SA, ethylene, ABA, and auxin treatment. In parallel studies, a correlation was observed between multiple cannabinoid biosynthetic pathway genes and SA application to the cannabis growing medium, along with a correlation between MeSA floral application and an increase in cannabinoid content. The results from all aspects of this study demonstrated an interaction between certain hormones and cannabinoid synthesis.

## Introduction

*Cannabis sativa*, an herbaceous flowering plant, is emerging as a key player in the future of medicine. Along with terpenes, the production of cannabinoids by the translation products of the cannabinoid biosynthetic pathway genes (and factors that affect their expression) determine the medicinal value of the plant. The bulk of cannabinoid production occurs predominantly in unfertilized female cannabis flowers and increases substantially as these female flowers undergo developmental changes during maturation. There is a lack of understanding on how these genes are regulated throughout the flowering period, despite the potential importance of this regulation in terms of the impact that cannabinoid production has on the value of this crop plant.

The cannabinoid biosynthetic pathway is an intricate combination of various pathways, with many products including terpenes, flavonoids, sterols, and ultimately, they form a pathway that produces cannabinoids (Andre et al. 2016). There are over 100 identified cannabinoids, which are naturally occurring compounds in cannabis plants that interact with the endocannabinoid system in the human body (Gertsch et al. 2010).

The precursors of cannabinoids originate from two pathways: the polyketide pathway and the plastidial 2-C-methyl-D-erythritol 4-phosphate (MEP) pathway (Blatt-Janmaat and Qu 2021). The polyketide pathway gives rise to olivetolic acid (OA), while the MEP pathway gives rise to geranyl diphosphate (GPP). The MEP pathway both develops monoterpenes and initiates cannabinoid production. GPP is the precursor to monoterpenes, and it is involved in the reaction which produces cannabigerolic acid (CBGA). Importantly, how the bifurcation of GPP as a precursor to monoterpenes and cannabinoids is regulated is completely unknown. Aromatic prenyltransferase (AP) enzymes catalyze the alkylation of GPP and OA, to form CBGA. CBGA is the precursor to many end-point cannabinoids, a few key ones including cannabidiolic acid (CBDA), tetrahydrocannabinolic acid (THCA), and cannabichromenic acid (CBCA) (Blatt-Janmaat and Qu 2021).

Previous studies have focused on the cannabinoid synthase gene, tetrahydrocannabinolic acid synthase (THCAS), as the rate-limiting enzyme in the production of THCA, likewise with CBDAS and CBDA (Muntendam et al. 2009; Richins et al. 2018). However, conclusions from more recent studies have determined that the correlation between THCAS gene expression and THCA content is poor, and attention should be paid to other enzymes in the cannabinoid biosynthetic pathway (Cascini et al., 2013, 2019; Y. Liu et al., 2021). In recent work from this lab, this point was clearly established and further, the production of CBGA was identified as rate-limiting end point cannabinoid synthesis (Apicella et al. 2022).

Research indicates that geranyl diphosphate synthase (GPPS) rate-limits the production of monoterpenes in various plant species, such as orchids (Chuang 2017). Monoterpene biosynthesis in plants is largely conserved, indicating that GPPS is likely rate-limiting in monoterpene production in cannabis, while simultaneously being involved in cannabinoid synthesis. The AP enzymes utilize GPPS’s product, geranyl diphosphate (GPP), to produce CBGA. This recent research has led our team to believe that studying the genes that facilitate CBGA production will lead us to understanding cannabinoid production at a more detailed level.

There are currently two identified aromatic prenyltransferases that are capable of producing CBGA in cannabis, prenyltransferase 1 (CsPT1) and prenyltransferase 4 (CsPT4). They can catalyze the alkylation of OA and GPP to form CBGA. Aromatic prenyltransferases are membrane-bound enzymes, localized to plant plastids, (de Bruijn et al. 2020). Both CsPT1 and CsPT4 are known to produce CBGA, however various studies have shown that CsPT4 produces more CBGA and has a higher affinity for its substrate, OA (Blatt-Janmaat and Qu 2021).

CsPT1 and CsPT4 are the focus of this research, along with the molecular signals which initiate trichome development and cannabinoid production. These processes depend on hormonal signals, which induce transcriptional activators and repressors (Fambrini and Pugliesi 2019). There are many hormones involved in plant regulation and development. We focus on the major ones including salicylic acid (SA) (and methyl salicylate (MeSA)), gibberellic acid (GA), auxin (NAA), abscisic acid (ABA), cytokinin (trans-Zeatin), ethylene (ACC), jasmonic acid (JA) (and methyl jasmonate (MeJA)).

Hormones can work synergistically, antagonistically, or in solitude (Wang and Irving 2011; Lehmann et al. 2020). Very few studies have been published regarding how hormone signaling pathways affect cannabis plants. The published research takes time to emphasize the important role of hormones in cannabinoid production (Mansouri et al. 2009; Mansouri et al. 2013; Jalali et al. 2019; Apicella et al. 2022). For example, in (Apicella et al. 2022), it is shown that jasmonic acid signaling pathways can increase cannabinoid content in flowering plants.

Our research encompasses a variety of assays conducted on *Cannabis sativa* and its key enzymes. CsPT1 and CsPT4 (putative GOT enzymes) are examined in both their promoter regions and their coding domain regions. Promoter regions are the upstream sequences prior to the transcription start site of the gene of interest (Hernandez-Garcia and Finer 2014). These regions contain responsive elements, identified through online bioinformatic analyses, which are binding sites for various transcription factors that are activated through signaling pathways (Higo et al. 1999). The goal of this research is to understand how key, potentially rate-limiting, genes are regulated through hormone networks. This understanding will aid to figuring out how processes are occurring during female flower development and why cannabinoid production occurs.

## Results

### Expression patterns of *CsPT1* and *CsPT4*

Real-time PCR was conducted on different tissues in the female cannabis plant, and root expression was used as the control. Results showed that both *CsPT1* and *CsPT4* are highly expressed in the flowers, and they are also present in the leaves, indicating their primary function in the flower tissue (Figure 1). We further analyzed GUS activity driven by the *CsPT1* and *CsPT4* promoters (*CsPT1 pro* and *CsPT4 pro*). It was found that the activities of *CsPT1 pro* and *CsPT4 pro* were observed in the young leaves of cannabis seedlings (Figure 1b). In addition, the activity of *CsPT4 pro* was much stronger toward the base of the sugar leaves, in which area the capitate stalked glandular trichomes are predominantly developed (Figure 1c), further indicating the main function of *CsPT4* in these glandular trichomes.

**Fig. 1.**
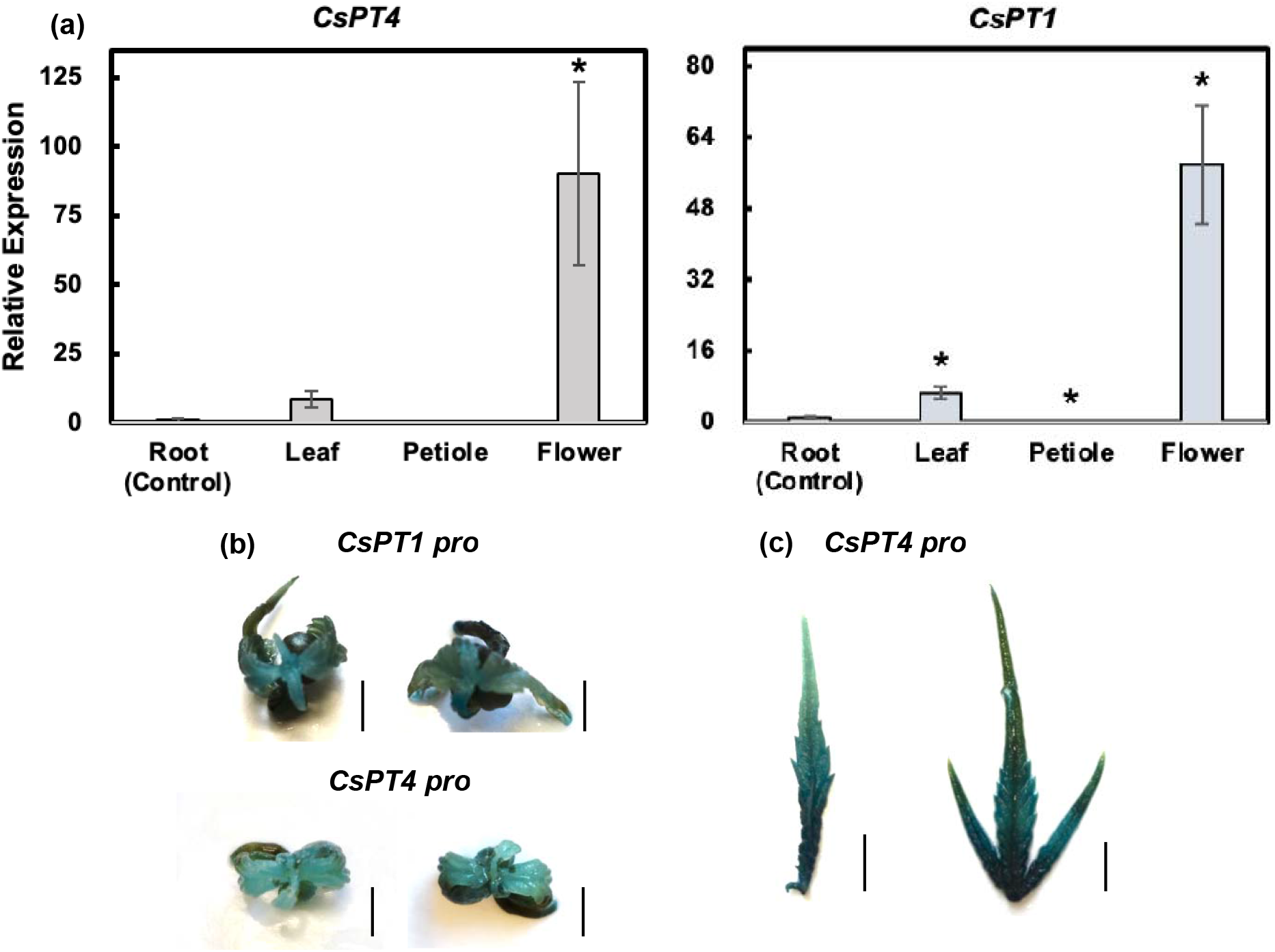
*CsPT4* and *CsPT1* expression pattern in cannabis tissues. a Transcript levels of *CsPT4* and *CsPT1* in various tissues, including root, leaf, petiole, and flower. Expression levels of each tissue were compared to the root. Student’s *t*-test was utilized for calculating statistical significance, * indicates *p* ≤ 0.05 (n=3). b, c Transient transfection of *CsPT4 pro* and *CsPT1 pro* driven GUS reporter in cannabis seedlings (b) and sugar leaves (c). Scale bars represent 5 mm.

### *In silico* analysis of *CsPT1 and CsPT4* promoters

Regarding *CsPT4 pro*, there is likely more sequences contained in the promoter region, upstream of the section we were able to clone, with additional responsive elements. However, upstream of the 1500 bp promoter region of *CsPT1* lies a coding domain region for another gene, indicating that this is the entire promoter region. As shown in Figure 1b, 600 bp of *CsPT4 pro* is sufficient to have a strong activity. Analysis using PLACE and PlantCARE, the online query software for cis-acting regulatory element analyses, revealed many putative responsive elements in the cloned promoter fragments of *CsPT1* and *CsPT4* (Higo et al. 1999). These elements are conserved sequences which have been demonstrated, in other organisms, to bind to transcription factors related to various environmental cues such as stress, wounding, drought, and hormone signaling.

The hormone responsive elements for both *CsPT1 pro* and *CsPT4 pro* are listed in Table 1. The *in silico* analysis showed that in *CsPT4 pro*, there are three responsive elements for cytokinin, one for SA, one for GA, one for ethylene, and one for ABA. The analysis also revealed that in *CsPT1 pro*, there are five GA responsive elements, two SA, six auxin, and three ABA responsive elements. MeJA signaling is vital for glandular trichome formation and secondary metabolite biosynthesis (van der Fits and Memelink 2000; Li et al. 2015; Yan et al. 2017; Zhang et al. 2020; Hua et al. 2021; Yan et al. 2022). However, no JA responsive element is present in the two promoters, even in a larger *CsPT4 pro* region. It is possible that JA does not regulate transcription factors that activate *CsPT1* and *CsPT4*.

**Table 1.** *In silico* identification of hormone responsive elements of *CsPT1* and *CsPT4* promoters*.

| Promoter | *CsPT1* (1500 bp region) | *CsPT4* (600 bp region) |
| --- | --- | --- |
| Cytokinin | 0 | 3 |
| Gibberellin | 5 | 1 |
| Abscisic Acid | 3 | 1 |
| Salicylic Acid | 2 | 1 |
| Ethylene | 0 | 1 |
| Auxin | 3 | 0 |

\* The promoters were analyzed using PLACE and PlantCARE online programs.

### *CsPT4 pro* exhibits a strong response towards various hormones

The 600 bp region of *CsPT4 pro* that was cloned contains three putative cytokinin responsive elements; 2 kb *CsPT4 pro* was predicted to have no auxin responsive elements. For this reason, NAA was used as a negative control. 3 and 5 hours post treatment (hpt) from ZR (cytokinin), *CsPT4 pro* activated the luciferase reporter at significant levels (Figure 2a). The result supports the *in silico* analysis that cytokinin may regulate *CsPT4* expression. Because there are no predicted auxin responsive elements in *CsPT4 pro*, it is expected that there is no increased luciferase activity upon auxin treatment. Following application of NAA, there was no increased activity measured from *CsPT4 pro*, indicating that auxin may not modulate *CsPT4* transcription in Arabidopsis protoplasts (Figure 2b).

**Fig. 2.**
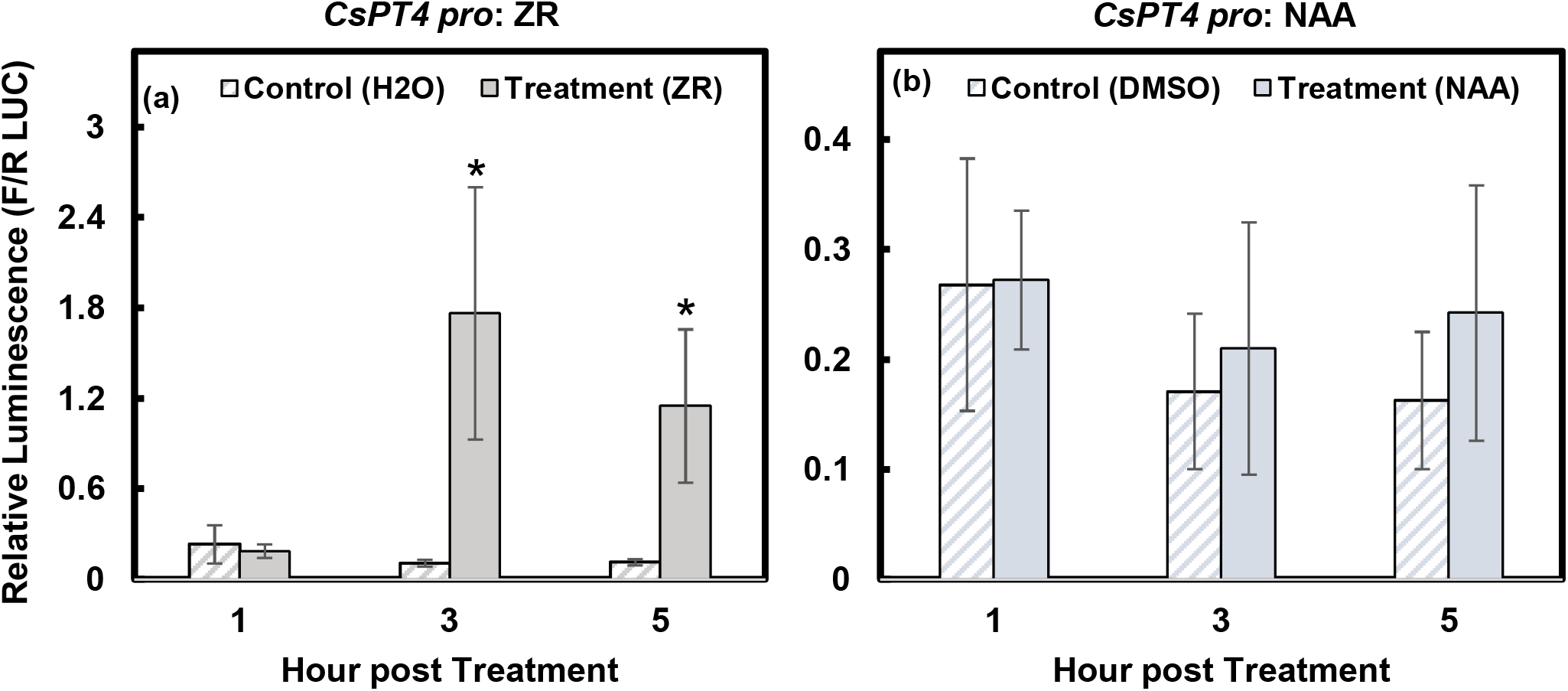
Responses of *CsPT4 pro* to ZR and NAA in Arabidopsis protoplasts. Dual luciferase assay results showed the activity of the *CsPT4 pro* indicated by relative luminescence, calculated by a ratio of firefly LUC to *Renilla* LUC. Responses to ZR (a) and NAA (b) at 1, 3, and 5 hpt are shown. A Student’s *t*-test was used to compare relative luminescence of treated protoplasts to the solvent control at each time point. This method of statistical analysis was repeated for each DLR assay (Figures 3-5). * indicates *p* ≤ 0.05 (n=3).

In addition, the 600 bp *CsPT4* promoter region contains one ethylene, one ABA, and one GA putative responsive element. At 5 hpt with ACC (ethylene), *CsPT4 pro* was able to be activated at significant levels (Figure 3a). In response to ABA and GA_3_, *CsPT4 pro* was activated at 3 and maintained at 5 hpt (Figure 3b,c). These three hormones had similar levels of *CsPT4 pro* activation. The results indicate that *CsPT4* can be involved in different hormonal signaling pathways; hormones may coordinate to control glandular trichome development and cannabinoid biosynthesis in cannabis female flowers.

**Fig. 3.**
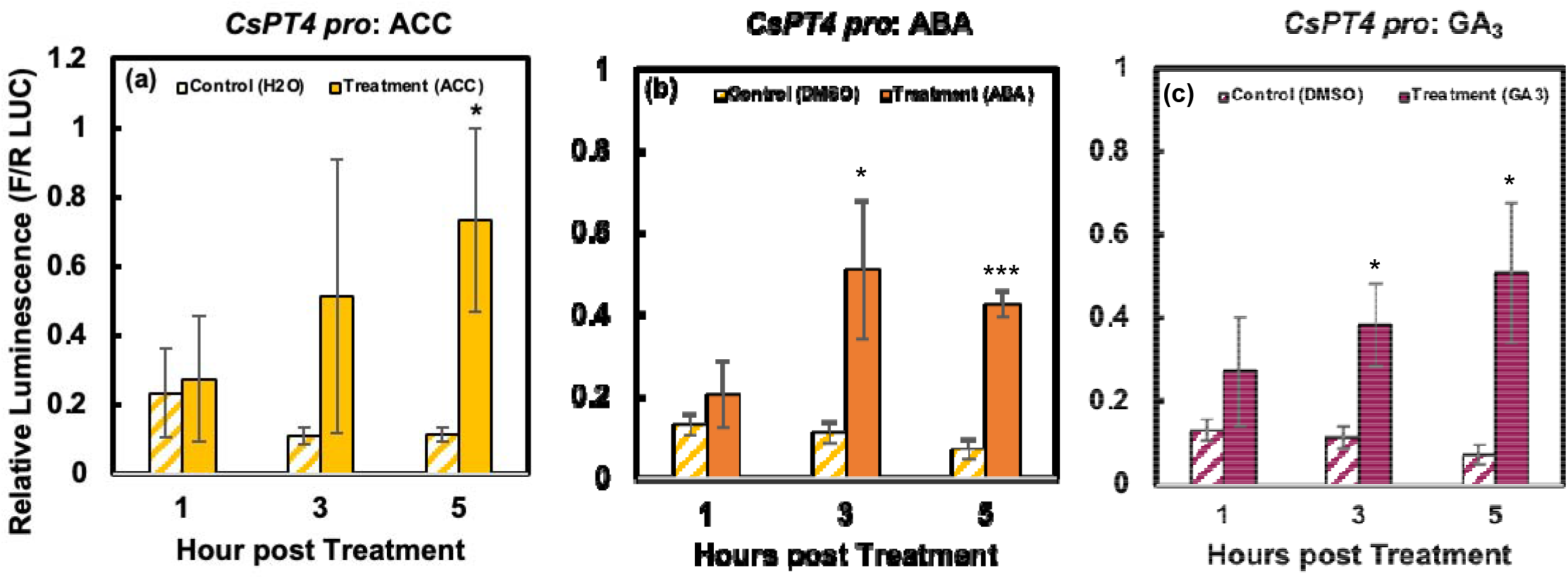
Responses of *CsPT4 pro* to ACC, ABA, and GA. Dual luciferase assay results display the activation of *CsPT4 pro* over 5 hours indicated by relative luminescence, calculated by a ratio of firefly to *Renilla* LUC. Significant activation seen from ACC (a), ABA (b), and GA_3_ (c). * indicates *p* ≤ 0.05. *** indicates *p* ≤ 0.001 (n=3).

### Auxin and ABA activate *CsPT1 pro*

The 1500 bp of *CsPT1 pro* contains three auxin, three ABA, and five GA putative responsive elements. Following treatment from NAA, *CsPT1 pro* was activated significantly as early as 1 hpt. The activity was slightly higher at 3 hpt and retained at 5 hpt (Figure 4a). ABA was able to induce significant activation of *CsPT1 pro* only at 3 hpt and the activity returned to the basal level 5 hpt (Figure 4b). Treatment with GA_3_ prompted slight increases in relative luminescence, however there was no significant difference compared to the solvent control at each time point (Figure 4c).

**Fig. 4.**
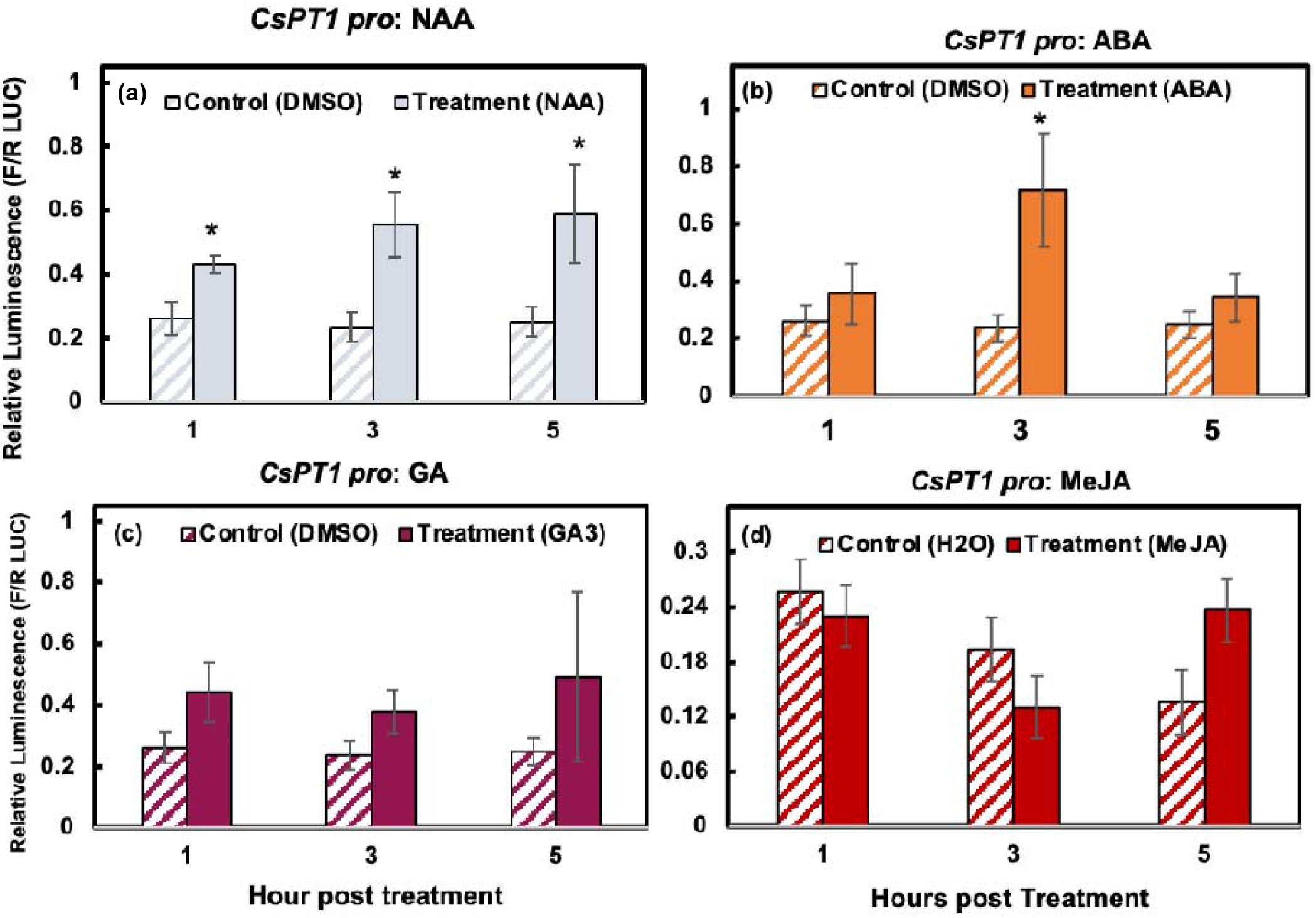
Responses of *CsPT1 pro* to ABA, NAA, GA, and MeJA. Dual luciferase assay results displaying activity of *CsPT1 pro* at 1, 3, and 5 hpt indicated by relative luminescence. Significant activation was measured from NAA (a) as well as ABA (b). No activation of *CsPT1 pro* was detected in response to GA_3_ (c) or MeJA (d). * indicates *p* ≤ 0.05. ** indicates *p* ≤ 0.01 (n=3).

MeJA is a major hormone involved in plant defense responses to stress stimuli (Cheong and Choi 2003; Truman et al. 2007; Wasternack and Hause 2013). Our previous study also showed that MeJA foliar spray increased cannabinoid production (Apicella et al. 2022). Although no MeJA responsive elements was found in *CsPT1 pro*, the response of *CsPT1 pro* towards MeJA was still examined. Interestingly, there was slight decrease of luminescence 3 hpt and then slight increase of luminescence 5 hpt (Figure 4d). However, there was no significant difference observed at either time point. MeJA could regulate other cannabinoid biosynthetic genes or promote glandular trichome morphogenesis, resulting in higher cannabinoid accumulation.

### *CsPT1 pro* and *CsPT4 pro* are responsive to SA

*CsPT1 pro* and *CsPT4 pro* were examined collectively for their response to SA. Both promoters were activated after treatment with SA (Figure 5). *CsPT1 pro* was activated sooner at 1 hpt and the activity was retained at similar levels from 1 to 5 hpt (Figure 5a). *CsPT4 pro* was not activated until 3 hpt with a slight decrease at 5 hpt (Figure 5b). Intriguingly, the increased activity of *CsPT4 pro* was substantially higher than that of *CsPT1 pro* despite one more SARE in the *CsPT1 pro*, suggesting a greater demand of *CsPT4* transcripts after SA elicitation.

**Fig. 5.**
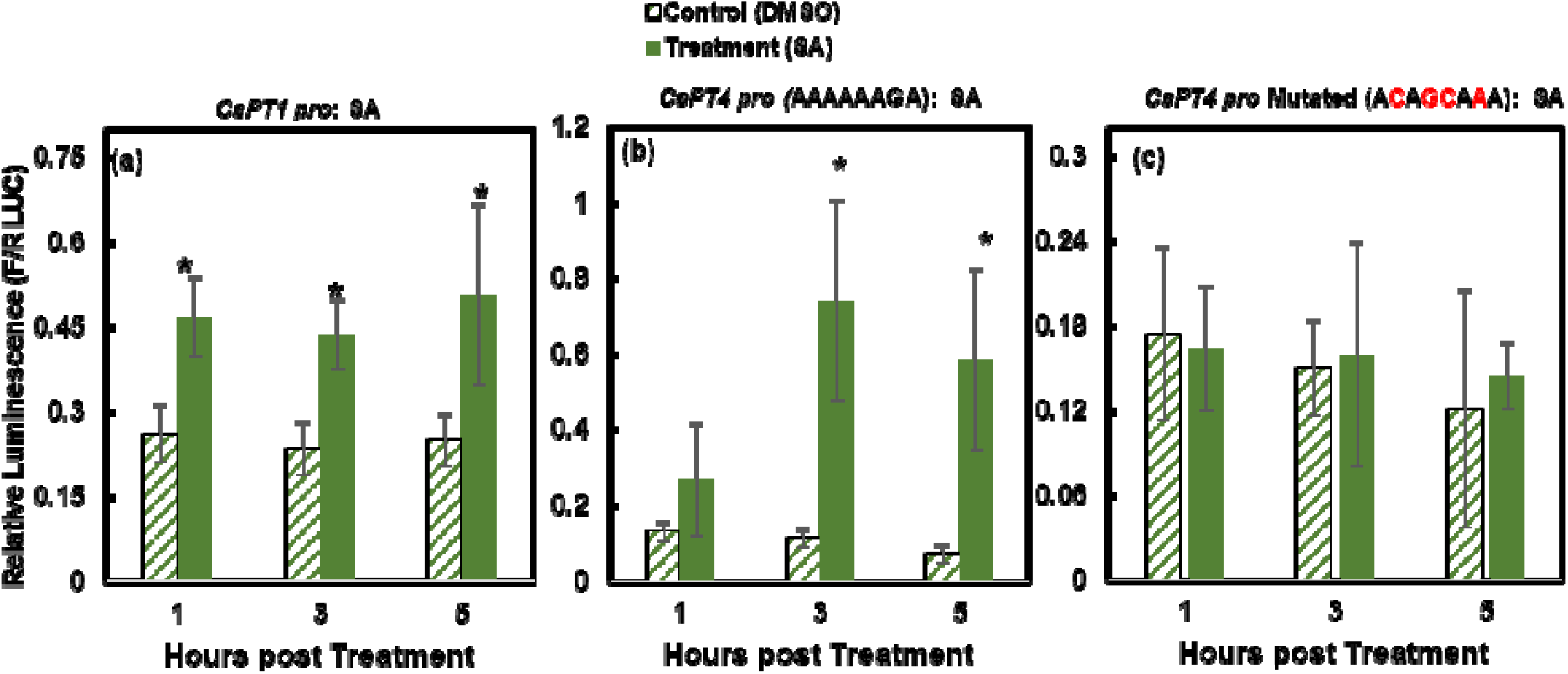
SA activated *CsPT1 pro* and *CsPT4 pro*. Dual luciferase assay results displaying activation of *CsPT1 pro* (a) and *CsPT4 pro* (b) across 5 hr indicated by relative luminescence. Mutated *CsPT4 pro* (c) did not respond to SA. * indicates *p* ≤ 0.05 (n=3).

To further understand whether the identified putative SARE in *CsPT4 pro* is truly functioning, site-directed mutagenesis (SDM) was performed to introduce mutations in the sequence. Four base pairs were mutated to diminish or eliminate responses of *CsPT4 pro* to SA elicitation (Figure 5c). The results revealed that the mutated *CsPT4 pro* was no longer responsive to SA elicitation (Figure 5c), indicating that the identified SARE in *CsPT4 pro* is a *bona fide cis*-acting element that is involved in SA signaling.

### SA induces certain cannabinoid biosynthetic pathway genes

The SA root drench administered to flowering cannabis plants resulted in altered gene expression of six genes in the cannabinoid biosynthetic pathway. The genes examined include tetraketide synthase (*TKS*), olivetolic acid cyclase (*OAC*), geranyl diphosphate synthase small subunit (*GPPS ssu*), cannabidiolic acid synthase (*CBDAS*), and prenyltransferase 1 and 4 (*CsPT1* and *CsPT4*). We were able to identify predicted SARE in the promoters of *TKS, OAC, CBDAS, CsPT1*, and *CsPT4*.

*TKS* expression levels were significantly upregulated at every examined time point post treatment. *TKS* expression reached the peak 48 hpt and dropped 72 hpt but still significantly higher than the control (Figure 6a). *TKS* is a type III polyketide synthase and works with *OAC* to produce olivetolic acid. Interestingly, *OAC* transcript levels were reduced compared to time 0, however at each time point, the treated samples still showed higher expression levels compared to the control (Figure 6b).

**Fig. 6.**
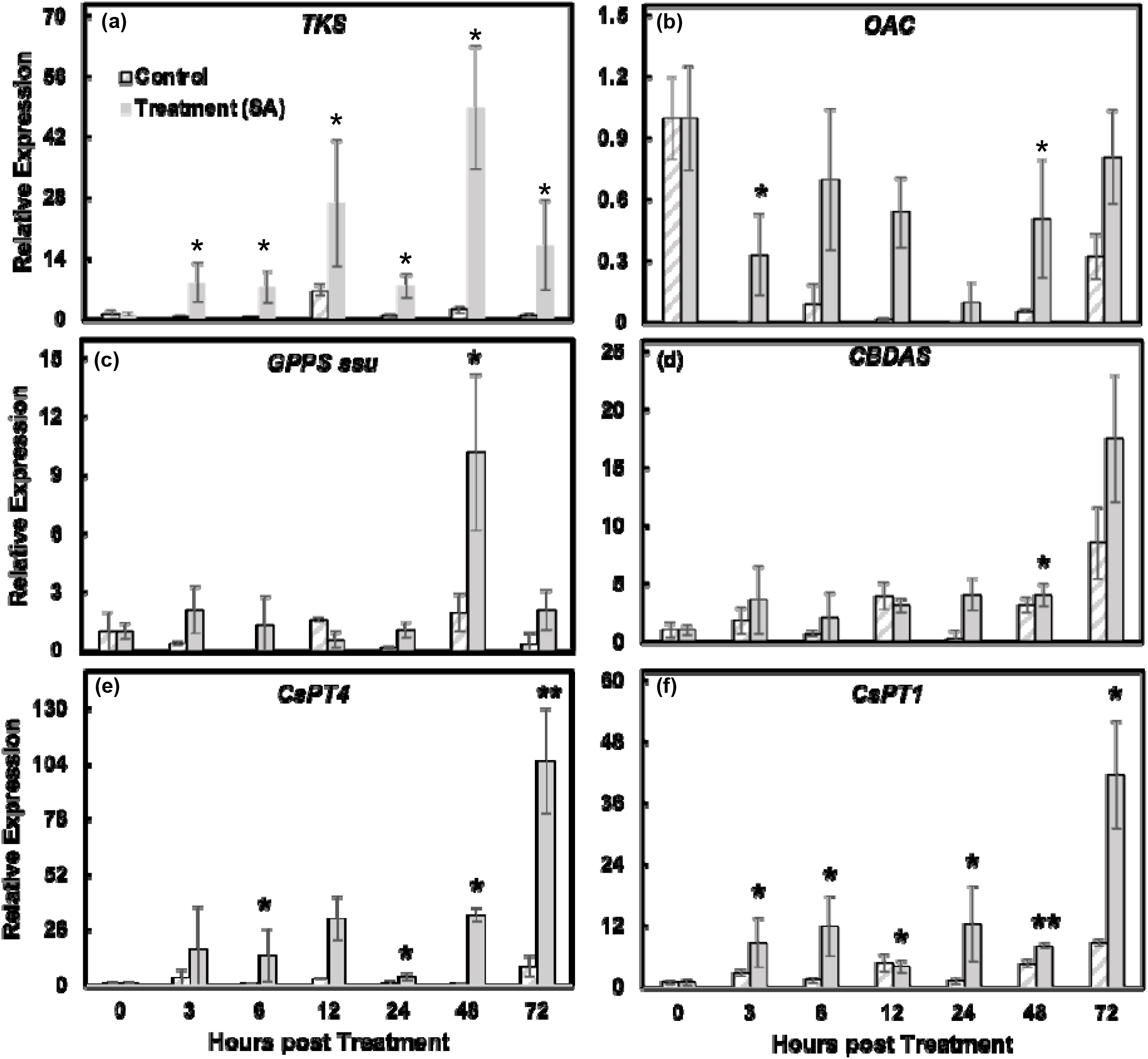
Real time PCR analysis of cannabinoid biosynthetic genes following SA treatment over a 72-hr period. Graphs shown include genes: (a) *TKS*, (b) *OAC*, (c) *GPPS ssu*, (d) *CBDAS* (e) *CsPT4* and (f) *CsPT1*. Results were averaged from four biological replicates. Statistical analysis was performed in comparison to DMSO control at each time point using Student’s *t*-test. * indicates *p* ≤ 0.05. ** indicates *p* ≤ 0.01. *** indicates *p* ≤ 0.001 (n=4).

It has been reported that the ratio of the small to large subunit of GPPS determines the amount of GPP synthesized (Orlova et al. 2009; Booth et al. 2017), therefore *GPPS ssu* was chosen for the analysis. *GPPS ssu* expression did not significantly increase until 48 hpt and then drastically dropped at 72 hpt (Figure 6c). In general, *CBDAS* was not upregulated by SA except for 24 hpt, which was mainly due to the decreased expression in the control samples; the expression level was similar to other time points except for 72 hpt (Figure 6d), suggesting that SA may not be responsible for inducing *CBDAS* expression. Interestingly, *CBDAS* transcripts increased 72 hpt compared to earlier time points in the control samples. SA increased *CBDAS* expression to a higher level at 72 hpt compared to the control, but no significant difference was observed (Figure 6d).

The *CsPT* gene expression levels were also induced by SA. Figure 6e showed that both *CsPT1* and *CsPT4* transcripts started to increase 3 or 6 hpt and were maintained at similar levels during the first 48 hpt, except for *CsPT4* at 24 hpt and *CsPT1* at 12 hpt. At 72 hpt, both genes increased substantially with *CsPT4* being unregulated around 100 folds compared to time the control. The gene expression analysis is consistent with results obtained from DLR assay (Figure 5a,b). Both CsPT4 and CsPT1 have been reported to be capable of synthesizing CBGA (Gülck et al. 2020; Luo et al. 2019; Jonathan E Page and Boubakir 2012), but the more recent findings showed that CsPT4 is the only enzyme that makes CBGA; CsPT1 failed to generate CBGA in *in vitro* assays in yeast and *Nicotiana benthamiana* (Gülck et al. 2020; Luo et al. 2019). The high expression level of *CsPT1* in the flowers and the upregulation by SA suggests that CsPT1 could be involved in the biosynthesis of cannabinoid under certain circumstances or the biosynthesis of other secondary metabolites in the glandular trichomes. Because all the assays were conducted in other organisms, it is also possible that CsPT1 functions differently in cannabis plants.

### MeSA increases cannabinoid biosynthesis

Because SA can induce cannabinoid biosynthetic genes (Figure 6), exogenous application of SA may potentially increase cannabinoid production. We showed that the foliar application of MeSA given to flowering plants resulted in a 47% increase in CBDA content per total flower weight (Figure 7). Statistical analysis did not find a significant difference between SA treated plants and DMSO treated control plants, likely due to high variability between replicates. However, we do see a trend of increased CBDA content. Future studies involving more biological replicates and more cannabinoid measurements, will likely create significant increases in cannabinoid content following SA treatments.

**Fig. 7.**
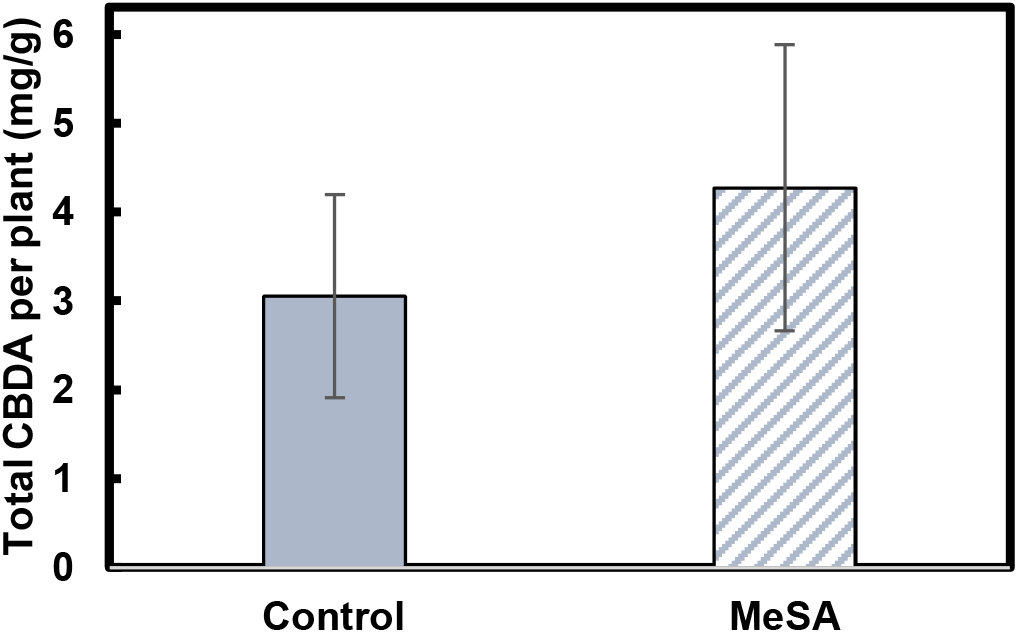
CBDA measurement following MeSA treatment. Slight CBDA increase was observed in treated plants compared to DMSO control plants (n = 3).

## Discussion

### CsPT1 vs. CsPT4 – Which enzyme plays a greater role for CBGA synthesis?

Both CsPT4 and CsPT1 have been shown to be able to produce CBGA from OA (Page and Boubakir 2012; Luo et al. 2019; Gülck et al. 2020). OA is the only substrate for CsPT4, and no other substrates were identified if there is any. On the contrary, CsPT1 is promiscuous and has numerous substrates including OA, olivetol, naringenin, and more (de Bruijn et al. 2020). *CsPT4* and *CsPT1* are both abundantly expressed in the flowers (Figure 1a). In addition, it was shown that the promoters of *CsPT1* and *CsPT4* are active in the leaves of young seedlings, with the stronger activities in the glandular trichomes. The data suggest that both genes should be evaluated as potential key enzymes facilitating cannabinoid production.

CsPT4 has shown greater enzymatic activity, with an affinity for OA at 6.73 μM (Luo et al. 2019), while the affinity of CsPT1 for OA was 60 mM (Page and Boubakir 2012), meaning that the affinity CsPT1 for the precursor of CBGA is 10000 times lower than that of CsPT4. Although both *CsPT1* and *CsPT4* are highly expressed in the flowers, the binding affinity implies that CsPT4 is more likely to be the enzyme that binds to OA for CBGA biosynthesis. In addition, when administered the necessary substrates in engineered yeast strains, CsPT4 produced much more CBGA (Luo et al. 2019). Our findings indicate that the promoters of *CsPT1* and *CsPT4* are responsive to different hormones, except for ABA and SA (Figures 2-5), suggesting that these two PT genes could be involved in discrete hormonal signaling pathways and hence, possibly responsible for different secondary metabolite biosynthesis. Previous research also showed that CsPT1 only showed enzymatic activity in *E. coli* but not in eucaryotic systems, such as yeast and *N. benthamiana* (Blatt-Janmaat and Qu 2021), suggesting functional differences between CsPT1 and CsPT4. While studying in other biological systems demonstrated some evidence of the functions of these two enzymes in the biosynthesis of CBGA, it is still not fully understood how the enzymes are acting in cannabis glandular trichomes. Further genetic analysis using *CsPT* mutants, techniques for overexpression, gene silencing or knock-out of the genes, will provide more explicit evidence on the involvement of these two CsPT enzymes in CBGA biosynthesis *in vivo*.

### Hormone regulation of *CsPT* promoters

Secondary metabolite biosynthesis is regulated by plant hormones (Zhang et al. 2020; Jan et al. 2021). Results from this research demonstrated that *CsPT4 pro* was responsive towards cytokinin, ABA, ethylene, SA, and GA, while *CsPT1 pro* was only activated following treatment with auxin, ABA, and SA (Figures 2-5), providing hints on the association between hormone signaling and cannabinoid biosynthesis.

ABA could activate both promoters in our study (Figures 3b, 4b). ABA plays essential roles in the regulation of floral development and plant responses to abiotic stress (Martignago et al. 2020; Hsu et al. 2021). Our previous study showed that both *CsPT1* and *CsPT4* were upregulated during female flower maturation and associated with glandular trichome development (Apicella et al. 2022). Previous studies demonstrated that ABA alone did not affect flower sex determination, however, ABA inhibited IAA induced female flower formation and impaired GA induced male flower development (Ram and Jaiswal 1972; Galoch 1978). ABA treatment increased THC concentration in both leaf and female flower tissues (Mansouri et al. 2009). The activation of the promoters of *CsPT1* and *CsPT4* by ABA suggest that ABA may also play important roles in glandular trichome development and cannabinoid biosynthesis during female flower maturation in cannabis plants.

The promoter of *CsPT4* was also activated by cytokinin, ethylene, and GA but not auxin (Figures 2 and 3). These hormones have been shown to be important for sex determination during floral development in dioecious or gynoecious plants (Galoch 1978; Huang et al. 2003; Wang et al. 2010; Ming et al. 2020; Punja and Holmes 2020; Adal et al. 2021). Cytokinin production in roots was shown to be essential for female sex determination in hemp plants generated from shoot cuttings of young plants (Chailakhyan and Khryanin 1978). Ethylene was shown to promote female flower development in industrial hemp (Galoch 1978; Mohan Ram and Sett 1982b; Mohan Ram and Sett 1982a; Moon 2020). Exogenous GA application induced male flower formation on female cannabis plants, which was inhibited by ABA (Ram and Jaiswal 1972). GA signaling associated genes were shown to be upregulated in induced-male (on female flowers) and male flowers compared to female flowers (Adal et al. 2021). In our study, NAA did not induce *CsPT1* and *CsPT4* promoters (Figure 4a), auxin (IAA) was shown to be involved in the induction of female flowers (Galoch 1978). Promoters of *CsPT1* and *CsPT4* were activated by hormones important for both female and male flower induction, indicating that *CsPT1* and *CsPT4* could be active in both female and male flowers. The findings also imply that cannabinoid biosynthesis involves interaction of different hormonal pathways that are essential for female flower induction and development. Monitoring of hormone contents during cannabis flower development will enable us to further understand how these hormones function in cannabinoids or other secondary metabolites biosynthesis in cannabis plants.

The DLR assays for hormone regulation of *CsPT1 pro* and *CsPT4 pro* were conducted using Arabidopsis protoplasts. Although many transcription factors are conserved among different plant species, there are also many with distinct functions in certain species. The results we observed in Arabidopsis protoplasts may be inconsistent or even contradictory. It would be interesting and important to further investigate the functions of these hormones in cannabis plants. Our findings still provide insights and clues to the hormonal modulation of glandular trichome development and cannabinoid biosynthesis in cannabis female flowers.

### SA plays important roles in the cannabinoid biosynthetic pathway

SA is primarily involved in plant defense responses to abiotic and biotic stresses. It is also an efficient elicitor for secondary metabolite biosynthesis (Singh et al. 2018; Ali 2021; Lv et al. 2021). As a plant undergoes abiotic or biotic stress, SA is produced at higher concentrations, in turn activating defense signaling pathways (Sun et al. 2016). Cannabinoids are known to be toxic to cells and must be exported to the secretory cavity (Sirikantaramas et al. 2005; Morimoto et al. 2007), where they may act as a defense mechanism against herbivory and in some cases, fungi and bacteria (Tanney et al. 2021). Our previous work showed that MeJA can increase cannabinoid production (Apicella et al. 2022). However, no JA responsive element was identified in the *CsPT* genes (Table 1). Here, we focused on the effects of SA on the cannabinoid biosynthetic pathway. One SARE in the cloned 600 bp of *CsPT4* pro was identified. After searching a broader region of the promoter, one more SARE was also identified. The only SARE in the 600 bp region was able to respond to SA application, which was further confirmed that the 600 bp of *CsPT4* pro with a mutated SARE failed to be activated by SA (Figure 5b). Moreover, the expression of *CsPT4* was substantially elevated by root drench of SA (Figure 6e). In addition, SA also increased other cannabinoid biosynthetic genes including *TKS, CBDAS, CsGPPS ssu*, and *CsPT1* (Figure 6). The findings provide strong evidence that SA can positively regulate cannabinoid biosynthesis.

SA has been shown to be involved in flowering in various plant species, such as Arabidopsis (Martínez et al. 2004), *Lemna pauciostate* (duckweed) (Khurana and Cleland 1992), and *Oryza sativa* (rice) (Izawa 2021). SA was also demonstrated to increase glandular trichome size and density in several species (Pandey et al. 2021; Zaid et al. 2022). Results from exogenous MeSA treatment was consistent with our hypothesis that SA application increased CBDA content (Figure 7). These findings imply a positive correlation between SA signaling and cannabinoid biosynthesis, which is likely rate-limited by CsPT4 and CBGA production.

### Experimental procedures

#### Plant materials and growth conditions

Wild type (Col-0) Arabidopsis seeds were surface sterilized with bleach, then plated on agar with half-strength Murashige and Skoog (MS) salts (Caisson Labs, Logan, UT, USA), MES (adjusted to pH 5.7 with Tris), 1% sucrose, and 0.8% agar, and grown in a growth chamber with a day (80–100 μmol m−2 s−1 illumination)/night cycle of 16h/8h at 22 °C. 8 days after germination, then seedlings were transferred into potting mix and grown in a growth chamber with LED lights at 12h light/12h dark at 22 °C.

Cannabis seed sterilization and seedling growth was conducted according to Sorokin et al. (2020). After germination, seedlings were placed on full MS (pH 5.6) plates for 10 more day. Then the seedlings were used for transient expression. Leaves were taken from flowering Auto Tsunami plants, an auto-flowering variety, grown in a growth chamber with LED lights of 16 h light/8 h dark at 26 °C. Agrobacterium mediated transient gene expression in cannabis seedlings and leaves followed the method published by Deguchi et al. (2020) with slight modifications.

Samples, Figure 1: Cuttings of 5-month-old Gorilla Glue (GG) mother plants were taken and rooted in an EZ-Cloner Classic ChamberTM (Sacramento, California) under aeroponic conditions. Rooted cuttings were grown under 18-h light/6-h darkness for 9 weeks in the closed-loop commercial facility, CTPharma LLC. Plants were then grown under 12h light/12h dark for reproductive growth.

Samples, Figure 7 & 8: Cuttings of a 5-month-old Wife mother plant, grown at the University of Connecticut, were taken, and rooted in Clonex rooting hormone (Hydrodynamics, MI, USA). Cuttings were grown under 18h light/6h dark. They were transplanted with Pro-Mix (AK, USA) Soilless mix. After 4 weeks in vegetative light, plants were grown under 12h light/12h dark for reproductive growth.

#### Cloning of promoters and *in silico* promoter analysis

Promoters of *Cannabis sativa CsPT1 pro* and *CsPT4 pro* were identified using the reference genome information on NCBI. For *CsPT4 pro*, we were able to clone 600 bp of the promoter, while we were able to clone 1500 bp of the *CsPT1 pro*. The promoters were PCR amplified using iProof High Fidelity Polymerase (Bio-Rad, CA, USA). *CsPT1 pro* and *CsPT4 pro* were inserted into an entry vector pDONR221, and then cloned into a destination vector, either P2GWL7 (for luciferase assay) or pBGWFS7 (for GUS assay). The orientation of insertion is shown in Supplementary Figure 1.

After the final insertion, plasmid maxiprep was conducted, with Macherey Nagel’s NucleoBond Xtra Maxi EF kit (Duren, Germany). *CsPT1 pro* and *CsPT4 pro* were analyzed using plant cis-acting regulatory DNA element databases, PLACE and PlantCARE. The predicted hormone responsive elements of *CsPT4 pro* and *CsPT1 pro* are shown in Supplementary Figure 2 and 3 respectively.

#### GUS Assay

GUS staining was performed according to the protocol in Arabidopsis: A Laboratory Manual (Weigel and Glazebrook 2002). GUS stained seedlings and leaves were recorded using a Cannon digital camera.

### Arabidopsis protoplast isolation and transfection

Arabidopsis leaves were collected from wt plants of 3-4 weeks old. Two protocols were followed for protoplast isolation. The tape-sandwich method was followed directly to separate the lower epidermal layer of the leaves, exposing the mesophyll cells (Wu et al. 2009). The remaining leaves were used to enzymatically isolate the protoplasts following a protocol published by Yoo, Cho, and Sheen (2007).

### Hormone treatment of protoplasts and real-time quantitative PCR

Hormones were administered to Arabidopsis protoplasts at concentrations of 5, 10, and 50 μM, to determine the concentration that would be used in the dual luciferase reporter assay (DLR). These concentrations were shown to cause a response in Arabidopsis (Lehmann et al. 2020). Seven hormones were administered to protoplasts, obtained from Sigma-Aldrich (MO, USA): cytokinin (trans-zeatin riboside, ZR), auxin (1-naphthaleneacetic acid, NAA), gibberellic acid_3_ (GA), salicylic acid (SA), abscisic acid (ABA), ethylene (1-aminocyclopropane-1-carboxylic acid, an ethylene precursor, ACC), and methyl jasmonate (MeJA).

Representative hormone responsive genes were chosen for qPCR (Supplementary Table 1) to ensure the hormones were activating the signaling pathways required for DLR assays (Hutchison and Kieber 2002; Israelsson et al. 2004; Davière and Achard 2013; Ren and Gray 2015; Mao et al. 2016; Leyser 2018; Lehmann et al. 2020; Hsu et al. 2021). Protoplasts were treated with various concentrations of hormones, then collected after 4 hours of shaking at 40 rpm in the dark, for RNA isolation and qPCR analysis.

Total RNA was isolated using TriReagent (Molecular Research Center, OH, USA) following the manufacturer’s protocol. cDNA was synthesized using iScript™ Reverse Transcription Supermix (Bio-Rad). qPCR was performed using iTaq Universal SYBR Green Supermix (Bio-Rad) in a CFX Connect system, with AtGAPDH as the housekeeping gene. Primers are listed in Supplementary Table 3. The hormone concentration which caused the highest fold increase in the gene, relative to the control, was chosen for further promoter analysis and DLR (Supplementary Table 2).

### Dual Luciferase Reporter assay

Dual luciferase assays were conducted in which *CsPT1 pro* and *CsPT4 pro* were cloned upstream of the firefly LUC gene; 35S CaMV promoter driven *Renilla* LUC was used as control. Arabidopsis protoplasts were transfected with both constructs (10 ug firefly LUC plasmid, 10 ng *Renilla* LUC plasmid, McNabb et al. 2005). Relative luminescence is calculated by dividing the recorded firefly luminescence by the *Renilla* luminescence. An empty P2GWL7 plasmid was also transfected as a negative control, which ensured there was no background noise.

Following overnight transfection, the protoplasts were treated with hormones. *CsPT4 pro* received 10 μM ZR, 10 μM ACC, 50 μM SA, 50 μM GA, 10 μM NAA and 10 μM ABA. *CsPT1 pro* received 50 μM GA, 50 μM SA, 10 μM NAA, 10 μM MeJA and 10 μM ABA. Samples were collected at 1, 3, and 5 hours post treatment (hpt). ZR, SA, GA, and ABA were dissolved in DMSO, while ACC, NAA, and MeJA was dissolved in DI water. The control samples were administered equal proportions of the solvent.

DLR reporter assay system was used (Promega, WI, USA) with the only alteration being the lysis buffer. Cell culture lysis reagent (Promega) was used instead of passive lysis buffer. To analyze the samples, the protocol by Yoo et al (2007) was followed. CLARIOstar plus plate reader (BMG Labtech, Germany) was used. Statistical analysis was determined using a one tailed Student *t*-test, following Nieradka et al. (2014). The experiment was repeated three times.

### Site-directed mutagenesis

Site directed mutagenesis was conducted on the SA responsive element (SARE) in the cloned region of the *CsPT4 pro*. Q5 site directed mutagenesis kit (NEB, MA, USA) was used in which point mutations were made to the sequence of the putative SARE. See Supplementary Table 3 for primer design and Supplementary Figure 4 for sequence comparison. Transfection was repeated with the mutated promoter, and luminescence was measured following SA treatment as mentioned above.

### RNA isolation from cannabis tissues and qPCR

Approximately 250 mg of tissue was collected from the cannabis plants. Samples from GG included root, leaf, petiole, and flower. Samples from Wife were flowers. Tissues were flash frozen with liquid nitrogen and stored at -80 °C. Total RNA was isolated from samples using Plant and Fungi RNA Extraction Kit (Macherey Nagel, Duren, Germany). Single-stranded cDNA was synthesized as described above.

Primers for qPCR are listed in Supplementary Table 3. *CsPP2A* was used as the housekeeping gene for Wife samples, with 4 biological replicates for each treatment. *CsUbiquitin* was used for GG samples, with 3 biological replicates for each organ from the plant. Samples were analyzed using the delta delta ct method and an ANOVA was conducted using a paired Student’s *t*-test.

### SA treatment and HPLC analysis

Root Drench: SA was dissolved in 1 mL of DMSO and further diluted in 1 L of DI water, to obtain a 1 mM concentration. There were 4 biological replicates per treatment group. DMSO or SA was administered to the plants as a soil drench during the fourth week of flowering; treatment methods from other studies followed (Stevens et al. 2006; Press et al. 2007). Flower samples were collected at hour 0, 3, 6, 12, 24, 48, and 72 hours post treatment. Approximately 250 mg fresh flower samples were collected and flash frozen for qPCR assay.

Floral Spray: MeSA (Sigma-Aldrich) was dissolved in 1 mL of DMSO, then diluted in 1 L DI water to create a concentration of 100 μM. DMSO or MeSA was applied by foliar spray until flowers were fully soaked; treatment methods from other studies followed (Mayer 1997; Chen et al. 2003). The spray was applied during the fourth week of flowering, as a one-time treatment. Flower samples were collected from plants prior to treatment and two weeks post treatment. Cannabinoids were sampled by milling the entirety of each plant’s flowers into a composite mixture. Approximately 500 mg of flower tissue was used for cannabinoid extraction using 20 mL 9:1 methanol:dichloromethane (DCM). Samples were shaken for 1.5 hours, then disarded. Samples were then diluted with 100 μL extraction to 900 μL 9:1 methanol:DCM.

Cannabinoid concentration was analyzed on a Shimadzu High Pressure Liquid Chromatograph (HPLC) Instrument (LC-10). The column used was a Raptor Arc C18 (150 × 4.6 mm, 2.7 μm particle size) from Restek (PA, USA). The HPLC method used was Shimadzu’s protocol (Shimadzu 2020). Peaks on the chromatogram were identified using cannabinoids standards (Restek, PA).

## Supplementary Information

## Author Contribution

LBS, YM and GAB designed the research; LBS and SRH performed experiments and analyzed data; LBS, YM and GAB wrote the manuscript; GAB and YM supervised the research.

## Acknowledgment

We thank Frederick Pettit and Shelly Durocher for their assistant in maintaining cannabis and tobacco plants in the greenhouse. We thank Dr. Huanzhong Wang for providing us the p2GWL7 and pBGWFS7 vectors. This work was supported by Fine Fettle LLC. This work is supported by Foundational Knowledge of Plant Products AFRI 1028172 from the USDA National Institute of Food and Agriculture (YM and GAB), and Hatch project CONS01027 (GAB).

## References

Adal AM, Doshi K, Holbrook L, Mahmoud SS (2021) Comparative RNA-Seq analysis reveals genes associated with masculinization in female Cannabis sativa. Planta 253:1–17. https://doi.org/10.1007/S00425-020-03522-Y/FIGURES/6

Alazem M, Lin N-S (2017) Antiviral roles of abscisic acid in plants. Front Plant Sci 8

Ali B (2021) Salicylic acid: An efficient elicitor of secondary metabolite production in plants. Biocatal Agric Biotechnol 31:101884. https://doi.org/10.1016/J.BCAB.2020.101884

Andre CM, Hausman J-F, Guerriero G (2016) Cannabis sativa: The plant of the thousand and one molecules. Front Plant Sci 7. https://doi.org/10.3389/fpls.2016.00019

Apicella PV., Sands LB, Ma Y, Berkowitz GA (2022) Delineating genetic regulation of cannabinoid biosynthesis during female flower development in Cannabis sativa. Plant Direct 6:e412. https://doi.org/10.1002/PLD3.412

Blatt-Janmaat K, Qu Y (2021) The Biochemistry of Phytocannabinoids and Metabolic Engineering of Their Production in Heterologous Systems. Int J Mol Sci 22:2454. https://doi.org/10.3390/ijms22052454

Booth JK, Bohlmann J (2019) Terpenes in Cannabis sativa – From plant genome to humans. Plant Science 284:67–72

Booth JK, Page JE, Bohlmann J (2017) Terpene synthases from Cannabis sativa. PLoS One 12. https://doi.org/10.1371/journal.pone.0173911

Cascini F, Farcomeni A, Migliorini D, Baldassarri L, Boschi I, Martello S, Amaducci S, Lucini L, Bernardi J (2019) Highly predictive genetic markers distinguish drug-type from fiber-type cannabis sativa L. Plants 8. https://doi.org/10.3390/plants8110496

Cascini F, Passerotti S, Boschi I (2013) Analysis of THCA synthase gene expression in cannabis: A preliminary study by real-time quantitative PCR. Forensic Sci Int 231:208–212. https://doi.org/10.1016/j.forsciint.2013.05.019

Chailakhyan MKh, Khryanin VN (1978) The Role of Roots in Sex Expression in Hemp Plants

Chalvin C, Drevensek S, Dron M, Bendahmane A, Boualem A (2020) Genetic Control of Glandular Trichome Development. Trends Plant Sci 25:477–487

Chen F, D’Auria JC, Tholl D, Ross JR, Gershenzon J, Noel JP, Pichersky E (2003) An Arabidopsis thaliana gene for methylsalicylate biosynthesis, identified by a biochemical genomics approach, has a role in defense. Plant Journal 36:577–588. https://doi.org/10.1046/J.1365-313X.2003.01902.X

Chen Y-F, Etheridge N, Schaller GE (2005) Ethylene signaling pathway. Ann Bot 95:901–915

Cheong J-J, Choi Y do (2003) Methyl jasmonate as a vital substance in plants. Trends in Genetics 19:409–413. https://doi.org/10.1016/S0168-9525(03)00138-0

Chuang HW (2017) The dual function of harpin in disease resistance and growth in Phalaenopsis orchids. Orchid biotechnology III 309–329. https://doi.org/10.1142/9789813109223_0015

Coogan TA (2019) Analysis of the cannabinoid content of strains available in the New Jersey Medicinal Marijuana Program. J Cannabis Res 1. https://doi.org/10.1186/s42238-019-0011-z

Davière JM, Achard P (2013) Gibberellin signaling in plants. Development (Cambridge) 140:1147–1151. https://doi.org/10.1242/dev.087650

de Bruijn WJC, Levisson M, Beekwilder J, van Berkel WJH, Vincken JP (2020) Plant Aromatic Prenyltransferases: Tools for Microbial Cell Factories. Trends Biotechnol 38:917–934

Deguchi M, Bogush D, Weeden H, Spuhler Z, Potlakayala S, Kondo T, Zhang ZJ, Rudrabhatla S (2020) Establishment and optimization of a hemp (Cannabis sativa L.) agroinfiltration system for gene expression and silencing studies. Nature. https://doi.org/10.1038/s41598-020-60323-9

Deng H, Chen Y, Liu Z, Liu Z, Shu P, Wang R, Hao Y, Su D, Pirrello J, Liu Y, Li Z, Grierson D, Giovannoni JJ, Bouzayen M, Liu M (2022) SlERF.F12 modulates the transition to ripening in tomato fruit by recruiting the co-repressor TOPLESS and histone deacetylases to repress key ripening genes. Plant Cell 34:1250–1272. https://doi.org/10.1093/plcell/koac025

Fambrini M, Pugliesi C (2019) The dynamic genetic-hormonal regulatory network controlling the trichome development in leaves. Plants 8

Filgueiras CC, Martins AD, Pereira R v., Willett DS (2019) The ecology of salicylic acid signaling: Primary, secondary and tertiary effects with applications in agriculture. Int J Mol Sci 20

Galoch E (1978) The hormonal control of sex differentiation in dioecious plants of hemp (Cannabis sativd). The influence of plant growth regulators on sex expression in male and female plants. Acta Societatis Botanicorum Poloniae 47:153–162. https://doi.org/10.5586/asbp.1978.013

Gertsch J, Pertwee RG, di Marzo V (2010) Phytocannabinoids beyond the Cannabis plant - Do they exist? Br J Pharmacol 160:523–529

Gülck T, Booth JK, Carvalho Khakimov B, Crocoll C, Motawia MS, Møller BL, Bohlmann J, Gallage NJ (2020) Synthetic Biology of Cannabinoids and Cannabinoid Glucosides in Nicotiana benthamiana and Saccharomyces cerevisiae. J Nat Prod 83:2877–2893. https://doi.org/10.1021/ACS.JNATPROD.0C00241/ASSET/IMAGES/LARGE/NP0C00241_0009.JPEG

Hernandez-Garcia CM, Finer JJ (2014) Identification and validation of promoters and cis-acting regulatory elements. Plant Science 217–218:109–119

Higo K, Ugawa Y, Korenaga T (1999) Plant cis-acting regulatory DNA elements (PLACE) database: 1999

Hsu PK, Dubeaux G, Takahashi Y, Schroeder JI (2021) Signaling mechanisms in abscisic acid-mediated stomatal closure. Plant Journal 105:307–321. https://doi.org/10.1111/tpj.15067

Hua B, Chang J, Xu Z, Han X, Xu M, Yang M, Yang C, Ye Z, Wu S (2021) HOMEODOMAIN PROTEIN8 mediates jasmonateLJtriggered trichome elongation in tomato. New Phytologist 230:1063–1077. https://doi.org/10.1111/nph.17216

Huang S, Cerny RE, Qi Y, Bhat D, Aydt CM, Hanson DD, Malloy KP, Ness LA (2003) Transgenic Studies on the Involvement of Cytokinin and Gibberellin in Male Development. Plant Physiol 131:1270–1282. https://doi.org/10.1104/PP.102.018598

Hutchison CE, Kieber JJ (2002) Cytokinin signaling in Arabidopsis. Plant Cell 14. https://doi.org/10.1105/tpc.010444

Israelsson M, Mellerowicz E, Chono M, Gullberg J, Moritz T (2004) Cloning and overproduction of gibberellin 3-oxidase in hybrid aspen trees. Effects on gibberellin homeostasis and development. Plant Physiol 135:221–230. https://doi.org/10.1104/pp.104.038935

Izawa T (2021) What is going on with the hormonal control of flowering in plants? Plant Journal 105:431–445. https://doi.org/10.1111/tpj.15036

Jalali S, Salami SA, Sharifi M, Sohrabi S (2019) Signaling compounds elicit expression of key genes in cannabinoid pathway and related metabolites in cannabis. Ind Crops Prod 133:105–110. https://doi.org/10.1016/j.indcrop.2019.03.004

Jan R, Asaf S, Numan M, Lubna Kim KM (2021) Plant secondary metabolite biosynthesis and transcriptional regulation in response to biotic and abiotic stress conditions. Agronomy 11

Jeon HW, Park AR, Sung M, Kim N, Mannaa M, Han G, Kim J, Koo Y, Seo YS, Kim JC (2022) Systemic Acquired Resistance-Mediated Control of Pine Wilt Disease by Foliar Application With Methyl Salicylate. Front Plant Sci 12. https://doi.org/10.3389/fpls.2021.812414

Kalaivani K, Kalaiselvi MM, Senthil-Nathan S (2016) Effect of methyl salicylate (MeSA), an elicitor on growth, physiology and pathology of resistant and susceptible rice varieties. Sci Rep 6. https://doi.org/10.1038/srep34498

Khurana JP, Cleland CF (1992) Role of Salicylic Acid and Benzoic Acid in Flowering of a Photoperiod-Insensitive Strain, Lemna paucicostata LP6

Lehmann S, Dominguez-Ferreras A, Huang WJ, Denby K, Ntoukakis V, Schäfer P (2020) Novel markers for high-throughput protoplast-based analyses of phytohormone signaling. PLoS One 15. https://doi.org/10.1371/journal.pone.0234154

Leyser O (2018) Auxin signaling. Plant Physiol 176:465–479

Li A, Sun X, Liu L (2022) Action of Salicylic Acid on Plant Growth. Front Plant Sci 13. https://doi.org/10.3389/fpls.2022.878076

Li N, Han X, Feng D, Yuan D, Huang LJ (2019) Signaling crosstalk between salicylic acid and ethylene/Jasmonate in plant defense: Do we understand what they are whispering? Int J Mol Sci 20

Li S, Wang H, Li F, Chen Z, Li X, Zhu L, Wang G, Yu J, Huang D, Lang Z (2015) The maize transcription factor EREB58 mediates the jasmonate-induced production of sesquiterpene volatiles. The Plant Journal 84:296–308. https://doi.org/10.1111/TPJ.12994

Liu Y, Zhu P, Cai S, Haughn G, Page JE (2021) Three novel transcription factors involved in cannabinoid biosynthesis in Cannabis sativa L. Plant Mol Biol 106:49–65. https://doi.org/10.1007/s11103-021-01129-9

Liu Z, Ma C, Hou L, Wu X, Wang D, Zhang L, Liu P (2022) Exogenous SA Affects Rice Seed Germination under Salt Stress by Regulating Na+/K+ Balance and Endogenous GAs and ABA Homeostasis. Int J Mol Sci 23. https://doi.org/10.3390/ijms23063293

Livingston SJ, Quilichini TD, Booth JK, Wong DCJ, Rensing KH, Laflamme-Yonkman J, Castellarin SD, Bohlmann J, Page JE, Samuels AL (2020) Cannabis glandular trichomes alter morphology and metabolite content during flower maturation. Plant Journal 101:37–56. https://doi.org/10.1111/tpj.14516

Luján MA, Soria-García Á, Claver A, Lorente P, Rubio MC, Picorel R, Alfonso M (2021) Different Cis-Regulatory Elements Control the Tissue-Specific Contribution of Plastid ω-3 Desaturases to Wounding and Hormone Responses. Front Plant Sci 12. https://doi.org/10.3389/fpls.2021.727292

Luo X, Reiter MA, d’Espaux L, Wong J, Denby CM, Lechner A, Zhang Y, Grzybowski AT, Harth S, Lin W, Lee H, Yu C, Shin J, Deng K, Benites VT, Wang G, Baidoo EEK, Chen Y, Dev I, Petzold CJ, Keasling JD (2019) Complete biosynthesis of cannabinoids and their unnatural analogues in yeast. Nature 567:123–126. https://doi.org/10.1038/s41586-019-0978-9

Lv Z-Y, Sun W-J, Jiang R, Chen J-F, Ying X, Zhang L, Chen W-S (2021) Phytohormones jasmonic acid, salicylic acid, gibberellins, and abscisic acid are key mediators of plant secondary metabolites. World J Tradit Chin Med 7:307. https://doi.org/10.4103/WJTCM.WJTCM_20_21

Mansouri H, Asrar Z, Szopa J (2009) Effects of ABA on primary terpenoids and Δ9-tetrahydrocannabinol in Cannabis sativa L. at flowering stage. Plant Growth Regul 58:269–277. https://doi.org/10.1007/s10725-009-9375-y

Mansouri H, Salari F, Asrar Z (2013) Ethephon application stimulats cannabinoids and plastidic terpenoids production in Cannabis sativa at flowering stage. Ind Crops Prod 46:269–273. https://doi.org/10.1016/j.indcrop.2013.01.025

Mao JL, Miao ZQ, Wang Z, Yu LH, Cai XT, Xiang C bin (2016) Arabidopsis ERF1 Mediates Cross-Talk between Ethylene and Auxin Biosynthesis during Primary Root Elongation by Regulating ASA1 Expression. PLoS Genet 12. https://doi.org/10.1371/JOURNAL.PGEN.1005760

Martignago D, Siemiatkowska B, Lombardi A, Conti L (2020) Abscisic acid and flowering regulation: Many targets, different places. Int J Mol Sci 21:1–14

Martínez C, Pons E, Prats G, León J (2004) Salicylic acid regulates flowering time and links defence responses and reproductive development. Plant Journal 37:209–217. https://doi.org/10.1046/j.1365-313X.2003.01954.x

Mayer DF (1997) Effects of methyl salicylate on honey bee (Apis mellifera L.) foraging. N Z J Crop Hortic Sci 25:291–294. https://doi.org/10.1080/01140671.1997.9514018

McNabb DS, Reed R, Marciniak RA (2005) Dual luciferase assay system for rapid assessment of gene expression in Saccharomyces cerevisiae. Eukaryot Cell 4:1539–1549. https://doi.org/10.1128/EC.4.9.1539-1549.2005

Ming X, Tao Y bin, Fu Q, Tang M, He H, Chen MS, Pan BZ, Xu ZF (2020) Flower-specific overproduction of cytokinins altered flower development and sex expression in the perennial woody plant jatropha curcas l. Int J Mol Sci 21. https://doi.org/10.3390/ijms21020640

Mohan Ram HY, Sett R (1982a) Induction of fertile male flowers in genetically female Cannabis sativa plants by silver nitrate and silver thiosulphate anionic complex. Theoretical and Applied Genetics 1982 62:4 62:369–375. https://doi.org/10.1007/BF00275107

Mohan Ram HY, Sett R (1982b) Reversal of Ethephon-Induced Feminization in Male Plants of Cannabis sativa by Ethylene Antagonists. Zeitschrift für Pflanzenphysiologie 107:85–89. https://doi.org/10.1016/S0044-328X(11)80012-7

Moon Y-HYJSCMYCJ-KWT (2020) Effect of Timing of Ethephon Treatment on the Formation of Female Flowers and Seeds from Male Plant of Hemp (Cannabis sativa L.). Korean Journal of Plant Resources 33:682–688. https://doi.org/10.7732/KJPR.2020.33.6.682

Morimoto S, Tanaka Y, Sasaki K, Tanaka H, Fukamizu T, Shoyama Y, Shoyama Y, Taura F (2007) Identification and characterization of cannabinoids that induce cell death through mitochondrial permeability transition in cannabis leaf cells. Journal of Biological Chemistry 282:20739–20751. https://doi.org/10.1074/jbc.M700133200

Muntendam R, Erkelens T, Kayser O (2009) Genetic and Metabolic Studies of Cannabinoids in Standardized Medicinal Cannabis sativa. Planta Med 75. https://doi.org/10.1055/s-2009-1216452

Nieradka A, Ufer C, Thiadens K, Grech G, Horos R, van Coevorden-Hameete M, van den Akker E, Sofi S, Kuhn H, von Lindern M (2014) Grsf1-induced translation of the SNARE protein use1 is required for expansion of the erythroid compartment. PLoS One 9. https://doi.org/10.1371/journal.pone.0104631

Orlova I, Nagegowda DA, Kish CM, Gutensohn M, Maeda H, Varbanova M, Fridman E, Yamaguchi S, Hanada A, Kamiya Y, Krichevsky A, Citovsky V, Pichersky E, Dudareva N (2009) The small subunit of snapdragon geranyl diphosphate synthase modifies the chain length specificity of tobacco geranylgeranyl diphosphate synthase in planta. Plant Cell 21:4002–4017. https://doi.org/10.1105/tpc.109.071282

Page JE, Boubakir Z (2012) AROMATIC PRENYLTRANSFERASE FROM CANNABS

Page JE, Boubakir Z (2014) United States Patent Aromatic Prenyltransferase from Cannabis

Pandey N, Tiwari A, Rai SK, Pandey-Rai S (2021) Accumulation of Secondary Metabolites and Improved Size of Glandular Trichomes in Artemisia annua. 99–116. https://doi.org/10.1007/978-3-030-30185-9_31

Pokotylo I, Kravets V, Ruelland E (2019) Salicylic acid binding proteins (SABPs): The hidden forefront of salicylic acid signalling. Int J Mol Sci 20

Press CM, Wilson M, Tuzun S, Kloepper JW (2007) Salicylic Acid Produced by Serratia marcescens 90-166 Is Not the Primary Determinant of Induced Systemic Resistance in Cucumber or Tobacco. http://dx.doi.org/101094/MPMI1997106761 10:p761–768. https://doi.org/10.1094/MPMI.1997.10.6.761

Punja ZK, Holmes JE (2020) Hermaphroditism in Marijuana (Cannabis sativa L.) Inflorescences – Impact on Floral Morphology, Seed Formation, Progeny Sex Ratios, and Genetic Variation. Front Plant Sci 11:718. https://doi.org/10.3389/FPLS.2020.00718/BIBTEX

Ram HYM, Jaiswal VS (1972) Induction of male flowers on female plants of Cannabis sativa by gibberellins and its inhibition by abscisic acid. Planta 105:263–266. https://doi.org/10.1007/BF00385397

Ren H, Gray WM (2015) SAUR Proteins as Effectors of Hormonal and Environmental Signals in Plant Growth. Mol Plant 8:1153–1164

Richins RD, Rodriguez-Uribe L, Lowe K, Ferral R, O’Connell MA (2018) Accumulation of bioactive metabolites in cultivated medical Cannabis. PLoS One 13. https://doi.org/10.1371/journal.pone.0201119

Ruan J, Chen H, Zhu T, Yu Y, Lei Y, Yuan L, Liu J, Wang ZY, Kuang JF, Lu WJ, Huang S, Li C (2021) Brassinosteroids repress the seed maturation program during the seed-to-seedling transition. Plant Physiol 186:534–548. https://doi.org/10.1093/plphys/kiab089

Shimadzu (2020) The Potency Determination of 15 Cannabinoids using the Cannabis Analyzer for Potency™

Singh A, Dwivedi P, Padmanabh Dwivedi C (2018) Methyl-jasmonate and salicylic acid as potent elicitors for secondary metabolite production in medicinal plants: A review. J Pharmacogn Phytochem 7:750–757

Sirikantaramas S, Taura F, Tanaka Y, Ishikawa Y, Morimoto S, Shoyama Y (2005) Tetrahydrocannabinolic acid synthase, the enzyme controlling marijuana psychoactivity, is secreted into the storage cavity of the glandular trichomes. Plant Cell Physiol 46:1578–1582. https://doi.org/10.1093/pcp/pci166

Sorokin A, Yadav NS, Gaudet D, Kovalchuk I (2020) Transient expression of the β-glucuronidase gene in Cannabis sativa varieties. Plant Signal Behav 15. https://doi.org/10.1080/15592324.2020.1780037

Stevens J, Senaratna T, Sivasithamparam K (2006) Salicylic acid induces salinity tolerance in tomato(Lycopersicon esculentumcv. Roma): associated changes ingas exchange, water relations and membrane stabilisation. Plant Growth Regul. https://doi.org/10.1007/s10725-006-0019-1

Sun Y, Xia XL, Jiang JF, Chen SM, Chen FD, Lv GS (2016) Salicylic acid-induced changes in physiological parameters and genes of the flavonoid biosynthesis pathway in Artemisia vulgaris and Dendranthema nankingense during aphid feeding. Genetics and Molecular Research 15. https://doi.org/10.4238/gmr.15017546

Tahir MN, Shahbazi F, Rondeau-Gagné S, Trant JF (2021) The biosynthesis of the cannabinoids. J Cannabis Res 3. https://doi.org/10.1186/s42238-021-00062-4

Tanney CAS, Backer R, Geitmann A, Smith DL (2021) Cannabis Glandular Trichomes: A Cellular Metabolite Factory. Front Plant Sci 12

Truman W, Bennett MH, Kubigsteltig I, Turnbull C, Grant M (2007) Arabidopsis systemic immunity uses conserved defense signaling pathways and is mediated by jasmonates. Proc Natl Acad Sci U S A 104:1075–80. https://doi.org/10.1073/pnas.0605423104

van der Fits L, Memelink J (2000) ORCA3, a Jasmonate-Responsive Transcriptional Regulator of Plant Primary and Secondary Metabolism. Science (1979) 289:295–297. https://doi.org/10.1126/SCIENCE.289.5477.295

Wang D-H, Li F, Duan Q-H, Han T, Xu Z-H, Bai S-N (2010) Ethylene perception is involved in female cucumber flower development. The Plant Journal 61:862–872. https://doi.org/10.1111/j.1365-313X.2009.04114.x

Wang H, Zhao Q, Chen F, Wang M, Dixon RA (2011) NAC domain function and transcriptional control of a secondary cell wall master switch. Plant Journal 68:1104–1114. https://doi.org/10.1111/j.1365-313X.2011.04764.x

Wang YH, Irving HR (2011) Developing a model of plant hormone interactions. Plant Signal Behav 6:494–500. https://doi.org/10.4161/psb.6.4.14558

Wasternack C, Hause B (2013) Jasmonates: Biosynthesis, perception, signal transduction and action in plant stress response, growth and development. An update to the 2007 review in Annals of Botany. Ann Bot 111:1021–1058

Weigel Detlef, Glazebrook Jane (2002) Arabidopsis: A Laboratory Manual. Cold Spring Harbor Laboratory Press

Wu FH, Shen SC, Lee LY, Lee SH, Chan MT, Lin CS (2009) Tape-arabidopsis sandwich - A simpler arabidopsis protoplast isolation method. Plant Methods 5. https://doi.org/10.1186/1746-4811-5-16

Yan T, Chen M, Shen Q, Li L, Fu X, Pan Q, Tang Y, Shi P, Lv Z, Jiang W, Ma Y, Hao X, Sun X, Tang K (2017) HOMEODOMAIN PROTEIN 1 is required for jasmonate-mediated glandular trichome initiation in Artemisia annua. New Phytologist 213:1145–1155. https://doi.org/10.1111/nph.14205

Yan X, Cui L, Liu X, Cui Y, Wang Z, Zhang H, Chen L, Cui H (2022) NbJAZ3 is required for jasmonate-meditated glandular trichome development in Nicotiana benthamiana. Physiol Plant e13666. https://doi.org/10.1111/PPL.13666

Yoo SD, Cho YH, Sheen J (2007) Arabidopsis mesophyll protoplasts: A versatile cell system for transient gene expression analysis. Nat Protoc 2:1565–1572. https://doi.org/10.1038/nprot.2007.199

Yoon GM, Kieber JJ (2013) 1-Aminocyclopropane-1-carboxylic acid as a signaling molecule in plants. AoB Plants 5

Zaid A, Mohammad F, Siddique KHM (2022) Salicylic Acid Priming Regulates Stomatal Conductance, Trichome Density and Improves Cadmium Stress Tolerance in Mentha arvensis L. Front Plant Sci 13. https://doi.org/10.3389/fpls.2022.895427

Zhang R, Wang XJ, Gao W (2020) Regulation mechanism of plant hormones on secondary metabolites. China journal of Chinese materia medica 45:4205–4210. https://doi.org/10.19540/J.CNKI.CJCMM.20190129.007

