## Supplemental materials for "Analysis of hormonal regulation of promoter activities of *Cannabis sativa* prenyltransferase 1 and 4 and salicylic acid mediated regulation of cannabinoid biosynthesis"

Article Name:

Journal Name: Plant Molecular Biology,

Affiliation and email address of the corresponding author:

***CaMV 35S Pro***

***Renilla LUC***

**I. Control**

**Firefly LUC or GUS**

***CsPT1/CsPT4 Pro***

**II. Reporter**

**Supplementary** **Fig. 1**. Vectors used for DLR or GUS assays. I. Control vector contains *Renilla* LUC driven by the *CaMV* *35S* *pro*. II. Reporter vector contains *CsPT1 pro* or *CsPT4 pro* driven firefly LUC or GUS.

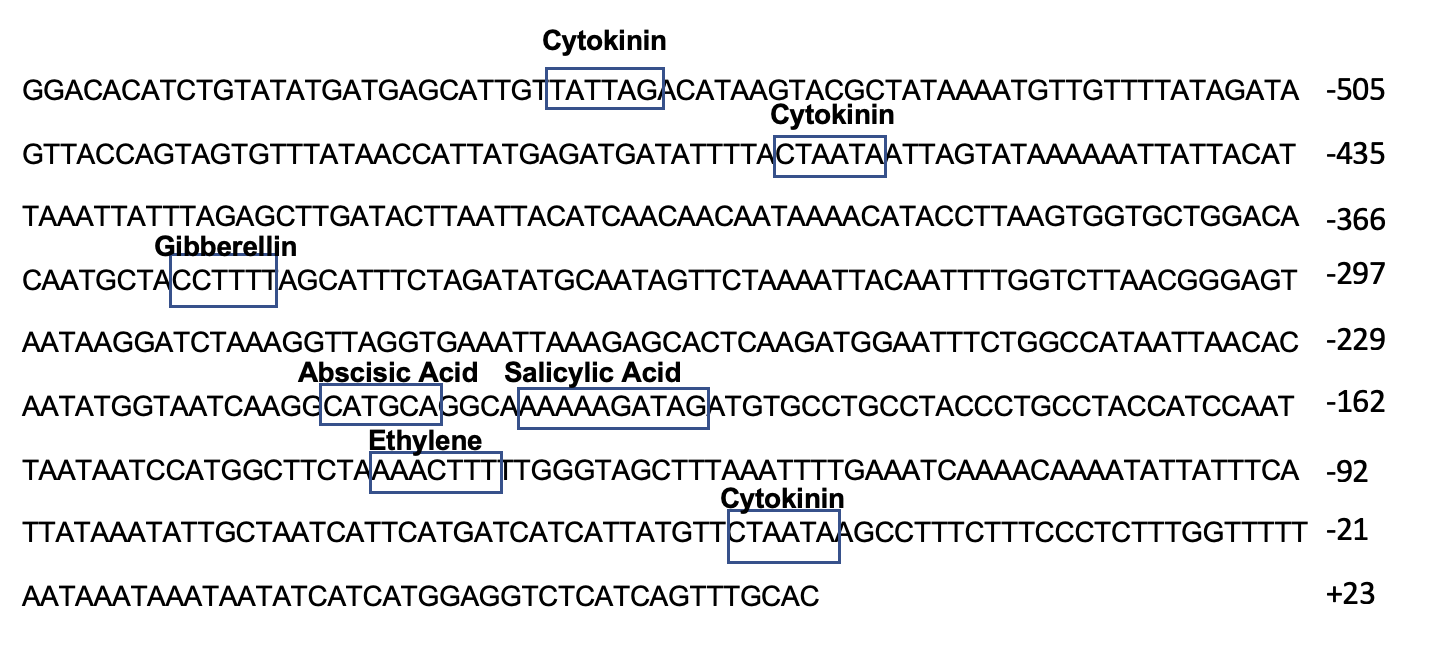

**Supplementary** **Fig. 2**. Hormone responsive elements within the 600 bp region of *CsPT4 pro* from an *in silico* analysis using PLACE and PlantCARE.

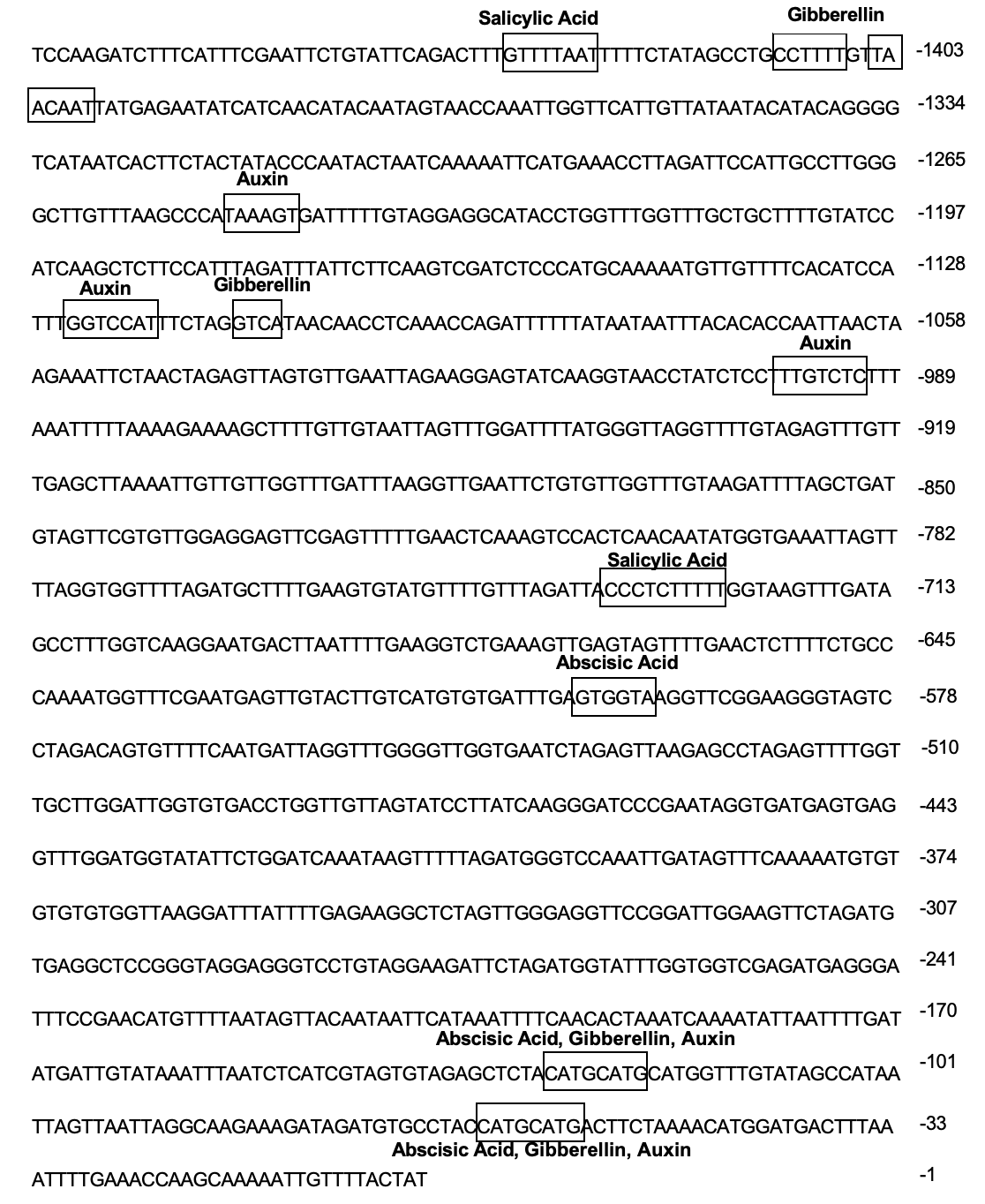

**Supplementary Fig. 3**. Hormone responsive elements within the 1500 bp region of *CsPT1 pro* from an *in silico* analysis from PLACE and PlantCARE.

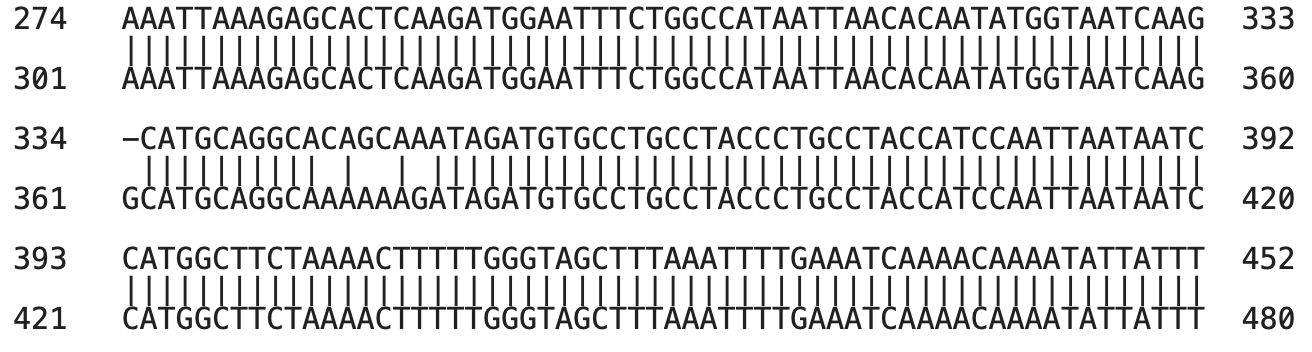

**SDM**

**NON-SDM**

**Supplementary** **Fig. 4.** Alignment of the original and site directed mutagenized SARE in *CsPT4 pro*. Four nucleotides are changed in the SARE, shown in red box and in Supplementary Table 3 in red font, to prevent binding from SA related transcription factors.

**Supplementary Table 1**. qPCR primers of Arabidopsis hormone responsive genes used in initial hormone experiment.

| Hormone Group | Gene | Forward Primer | Reverse Primer |
| --- | --- | --- | --- |
| Cytokinin | *AtARR2* | TCAGAGAACATCTTGCCTCGT | CGCGTTTTAGCTGCGAGTAG |
| Auxin | *AtSAUR41* | CCACCCGATCTTCGTTGGTT | GACCCTTCAAACCCTCCCAA |
| Gibberellic Acid_3_ | *AtGA3OX* | GGGGTGCCTTCCAAATCTCA | ACAGGTAGCCCGAAGAGACT |
| Salicylic Acid | *AtNPR1* | TCATTGCCGGAAGAGCTTGT | ACAGGTAGCCCGAAGAGACT |
| Abscisic Acid | *AtCOR78* | CGGGATTTGACGGAGAACCA | GGTCTCTTCCCAGCTCAGTC |
| Ethylene | *AtERF1* | CTAATCGAGCAGTCCACGCA | GTCCCGAGCCAAACCCTAAT |
| Methyl Jasmonate | *AtJAZ10* | TCCCATCGCAAGGAGAAAGT | AGGCCGATGTCGGATAGTAAG |
| Housekeeping Gene | *AtGAPDH* | CTCCCATGTTTGTTGTTGGTGTCA | CTTCCACCTCTCCAGTCCTTCATT |

**Supplementary Table 2**. qPCR of hormone responsive genes in response to various concentrations of their corresponding hormones

| **Hormone** | **Responsive Gene** | **Concentration (uM)** | **Fold Change**  **(relative to control)** |
| --- | --- | --- | --- |
| 1-Naphthaleneacetic Acid (Auxin) | Small Auxin Upregulated 41 (SAUR41) | 5 | 0.79 |
|  |  | 10 | 1.52 |
|  |  | 50 | 0.82 |
| Abscisic Acid | Cold Regulated 78 (COR78) | 5 | 0.88 |
|  |  | 10 | 2.72 |
|  |  | 50 | 2.42 |
| Salicylic Acid | Non-Expressor of Pathogenesis Related Genes 1 (NPR1) | 5 | 1.26 |
|  |  | 10 | 1.49 |
|  |  | 50 | 1.83 |
| Gibberellic Acid 3 | Gibberellin 3-Oxidase 1 (GA3OX) | 5 | 0.19 |
|  |  | 10 | 2.25 |
|  |  | 50 | 3.5 |
| Trans-Zeatin Riboside (Cytokinin) | Arabidopsis Response Regulator 2 (ARR2) | 5 | 0.41 |
|  |  | 10 | 1.08 |
|  |  | 50 | 0.91 |
| Methyl Jasmonate | Jasmonate-Zim-Domain Protein 10 (JAZ10) | 5 | 1.85 |
|  |  | 10 | 2.13 |
|  |  | 50 | 1.94 |
| 1-Aminocyclopropane-1-Carboxylic Acid (Ethylene) | Ethylene Response Factor 1 (ERF1) | 5 | 0.75 |
|  |  | 10 | 1.09 |
|  |  | 50 | 0.73 |

Note: Initial hormone experiment conducted in transfected Arabidopsis protoplasts. Multiple hormone concentrations were administered to the protoplasts. Highlighted in yellow is the gene expression level for the concentration to be used in DLR assays.

**Supplementary Table** **3.** Primers used for cloning, SDM and qPCR of cannabinoid biosynthetic genes (red nucleotides are the ones that were site-direct mutagenized).

| **Gene** | **FWD Primer** | **REV Primer** |
| --- | --- | --- |
| **qPCR primers** | | |
| ***CsPT1*** | TTGCTGGGATTATCTGGCCC | CTGCTTCCGGGTCGTAATTTG |
| ***CsPT4*** | CCATATCGAGTCATGTGGGCT | CCATATCGAGTCATGTGGGCT |
| ***CBDAS*** | CAGCAATTCCATTCCCTCAT | ATCCAGTTTAGATGCTTTTCGT |
| ***GPPS ssu*** | AGTTCACGAGGCCATGTACA | AAGGGAGGTGGTCATGAGTG |
| ***CsOAC*** | TCACAGAAGCCCAAAAGGAAGA | TACCCCTGCAATGGCAAACA |
| ***CsTKS*** | TCTTCACTAGCGCATCAACC | GAACGGTTCCACCACCATAA |
| ***CsUbiquitin*** | TACTGCGCCAGCTAACAAACC | GCACCCGTCTGACCTGAATC |
| ***CsPP2A*** | AGCAACGTTCAGCCCGTTAA | GACACTTCCCTCCAATTCGAAA |
| **Cloning and SDM primers** | | |
| ***CsPT4 promoter*** | GGACACATCTGTATATGATGAGC | GGACTCTCATTAGTTTGTACCTTTTC |
| ***CsPT1 promoter*** | TCCAAGATCTTTCATTTCGAATTC | ATAGTAAAACAATTTTTGCTTGGTTTC |
| **Attb1/2 sites** | GGGGACCACTTTGTACAAGAAAGCTGGGTC | GGGGACCACTTTGTACAAGAAAGCTGGGTC |
| ***CsPT4 promoter* SDM Primers** | CATGCAGGCACAGCAAATAGATGTGCCTGC | CTTGATTACCATATTGTGTTAATTATGGC  CAGAAATTCCATCTTG |
